# Biosynthesis of silver nanoparticles and exopolysaccharide using novel Thermophilic strain (Ts-1) of *Bacillus amyloliquefaciens*

**DOI:** 10.1101/2020.06.04.134742

**Authors:** Sharath Chandra Thota, Bathini Sreelatha

## Abstract

In the developing world, Nanotechnology became an efficient method in therapeutics, antimicrobials, diagnostics, catalysis, microelectronics, and high sensitivity biomolecular detection. As well as on the other hand, Exopolysaccharides are biopolymers which are also widely used in food formulation, bio- flocculants, bio-absorbents, drug delivery agents. As the chemical methods of synthesizing nanoparticles and polymers are environmentally risky, costly, and toxic. In the present study, we focused on production, purification, and characterization of silver nanoparticles (AgNPs) and exopolysaccharide (EPS) by eco-friendly, extracellular biosynthetic methods using novel thermophilic *Bacillus amyloliquefaciens* strain Ts-1. This strain was isolated from soil samples by employing pour and spread plate techniques. After obtaining pure culture, the bacterium was used for the synthesis of AgNPs and EPS. Nanoparticles were synthesized from AgNO_3_ by using reducing agents secreted by bacteria, and Exopolysaccharide biosynthesis is carried out in three steps by the organism in the presence of a carbon source. Synthesis of colloidal AgNPs and EPS was monitored by UV-Visible spectroscopy and Visual observation, respectively. SEM, Edax and FTIR were performed for the characterization of the AgNPs and EPS such as their size, morphology and composition and we also showed the catalytic activity of AgNps in degradation of methylene blue.

## Introduction

Nanoparticles are the particles with a size range of 1nm -100 nm. During the past few years, silver nanoparticles are among the most effective antimicrobial agents as mentioned in the original research article (Deljou.A et al. 2016) Nanoparticles are also widely used in diagnostics recently, gold nanoparticles are used in Detection of SARS-CoV-2 (Moitra et al. 2020). We know that physical and chemical techniques involved in synthesis of metal nanoparticles are very costly, toxic as well as not so efficient. Chemists and Biologists gathered to synthesis nanoparticles in an ecofriendly way of synthesizing metal nanoparticles using plants, bacteria, fungi, algae and yeast (Deljou.A et al. 2016) (Vilchis-Nestor et al. 2008)(Gholami-Shabani et al. 2014) (Govindaraju et al. 2008) (Kowshik et al. 2002).

Polysaccharides synthesized by bacteria were classified into three types based on their biological role. Like, glycogen - which is an intracellular polysaccharide that is stored inside the cell, K30 O-antigen - a polysaccharide which is capsular and closely linked to the cell surface, and cellulase - extracellular polysaccharides or exo-polysaccharides which are secreted out of the cells (wang et al. 2014)(Fontana et al. 2015). Biopolymers like EPS were not only produced by bacteria, also produced by plants, algae, fungi and yeast. (Gientka et al. 2015) (Dilna et al. 2015). Exopolysaccharides (EPSs) are heavy in their molecular weight and biodegradable (Sanalibaba et al. 2016). The production of EPS in bacteria carried out by four mechanisms which clearly described earlier by (Schmid et al. 2015). Application of EPSs is popular in various industries such as medicine, pharmaceuticals and cosmetics. Also, microbial EPSs can be used in the food industry (Bajpai et al. 2016) in some of the areas of usage are like controlling the viscosity and modify flow, Improve texture, mouthfeel, Thickeners softeners, Low calories food products, Dietary fibres, Films and coating agents, Frozen food icing, Moisturizing agents etc.

In the last two decades, The bacterial based product production has globally increased and got a good impact on their usage. Thus production is carried out in high quantities. In industries, they usually use large containers to make the desired product where temperature rises more than 40°C which is lethal to ‘mesophiles and psychrophiles’ which has to be controlled using high-level cooling machinery to cool the temperatures. To avoid these unwanted cooling systems which consume space, economy and skilled labour, many scientists were switching their focus on thermophiles that can withstand high temperatures and continue their metabolisms as usual without any lag. Many of Thermophilic enzymes are scientifically and industrially important. Thus exploration, identification and production of thermophiles and its products were increased.

Hence, In the present study, we are involved in the production of silver nanoparticles and exopolysaccharides from a thermophilic bacterium and also checked the degradation of methylene blue using AgNPs as a catalyst.

## Materials and Methods

### Media

The media used for enrichment and isolation of thermophilic bacteria is NAM (Nutrient agar media) and Emerson YpSs (Yeast Extract Soluble Starch) media. After trying with many other bacterial nutrient media, we got to know that YpSs broth is the best nutrient composition for this bacteria to produce EPS, and the Nutrient broth is good for the synthesis of AgNPs.

### Isolation and Identification

Soil samples were collected from five different places of Ramky Enclave’s litter dump yard “17.973517 N 79.610447 E” where temperature varies up to 40°-45°C. Samples were serially diluted and cultured using the pour plate method and spread plate method and incubated at 45°-50°C. The cultures were then sub-cultured on the NAM and YpSs agar plates for further studies using streaking. These colonies were observed for Gram’s nature, and preliminary identification was made using biochemical methods, as stated in Aneja’s manual. Further, the strain was identified by 16s rRNA sequencing.

### Synthesis of Silver nanoparticles

For the biosynthesis of silver nanoparticles, the selected bacterial isolate was inoculated into a 250-ml Erlenmeyer flask containing 100 ml sterile nutrient broth. The cultured flasks were incubated on a rotating shaker set at 200 rpm for 48 hrs at 45°c temperature. After incubation, the culture was centrifuged at 12,000 rpm for 10 min. The supernatant was collected and used for extracellular production of silver nanoparticles by mixing equal volume of filter-sterilized AgNO_3_ solution at 1000uM(1mM), 5000uM(5mM), 10000uM(10mM) of final concentrations. The reaction mixture was incubated on a rotating shaker (200 rpm) at room temperature for a period of 48hrs in the dark, and control also maintained (without adding AgNO3 solution to the cell-free supernatant). Visual observation was conducted periodically to check for the nanoparticle formation. Further characterization was performed for produced nanoparticles (Krishna et al. 2015).

### Extraction and Purification of EPS

Extraction and purification of EPS carried out in the same process described by (Aparna et al. 2017).

### Characterization of AgNps & EPS

The UV-visible spectrophotometer is used to observe the Plasmon absorbance in the range of 350-470 nm for the conformation of nanoparticle synthesis. Synthesized silver nanoparticles and exopolysaccharide size, shape and structural morphology are observed by Scanning Electron microscopy. EDAX is used to know the elemental composition of synthesized AgNps and EPS. The biotransformed products present in the extracellular filtrate were measured using Fourier Transform Infrared spectroscopy. For FTIR samples were freeze-dried and Diluted with potassium bromide in the ratio of 1: 100 and recorded with diffuse reflectance mode attachment in the range of 400-4000 cm-1 at a resolution of 4 cm-1.

### Catalytic application in dye degradation

Methylene blue is a heterocyclic aromatic chemical compound with molecular formula C_16_H_18_C_1_N_3_S. To investigate the catalytic activity of AgNPs, reduction of methylene blue was carried out using silver nanoparticles and incubating them for 48hrs. The extent of degradation of methylene blue dye using AgNPs as a catalyst was monitored by UV-Visible spectroscopy by measuring absorbance maxima of dyes at different time intervals 0h, 1h, 3h, 6h, 9h, 12h, 24h and 48h. A control set was maintained without AgNPs for dye and measured for absorbance. (Jyoti, K., & Singh, A. 2016).

## Results & Discussion

### Strain Isolation and Identification

The bacterium is identified by a series of biochemical tests and confirmed as gram-positive (Fig.1.b), Rod-shaped, endospore and pellicle forming (Fig.1.c) aerobic bacteria. As well the bacterium was found to be Catalase positive which can actively grow at 45°-50°C and can survive at 33° - 37°C (Fig.1.a.) According to the Aneja manual (Aneja et al. 2007), the biochemical results showed the present organism belongs to the genus *Bacillus*. Further, To find the species, it was sent for DNA sequencing which on the investigation of sequence through NCBI Blast Algorithm gave 98% homology to the *Bacillus amyloliquefaciens*, and this was submitted to the Genbank with a new strain name Thermophilic strain “Ts-1” with an accession number (MK553811.1). There are different strains of *Bacillus amyloliquefaciens* which are mesophiles; this is the first-ever report reporting *Bacillus amyloliquefaciens* strain Ts-1 as a thermophile that can grow above 45°C. Strain TS-1 was isolated and cultured accidentally on YePD media while isolating thermophilic fungi. We mistook it as yeast in the beginning till Grams and Biochemical tests were done.

**Fig. 1.**
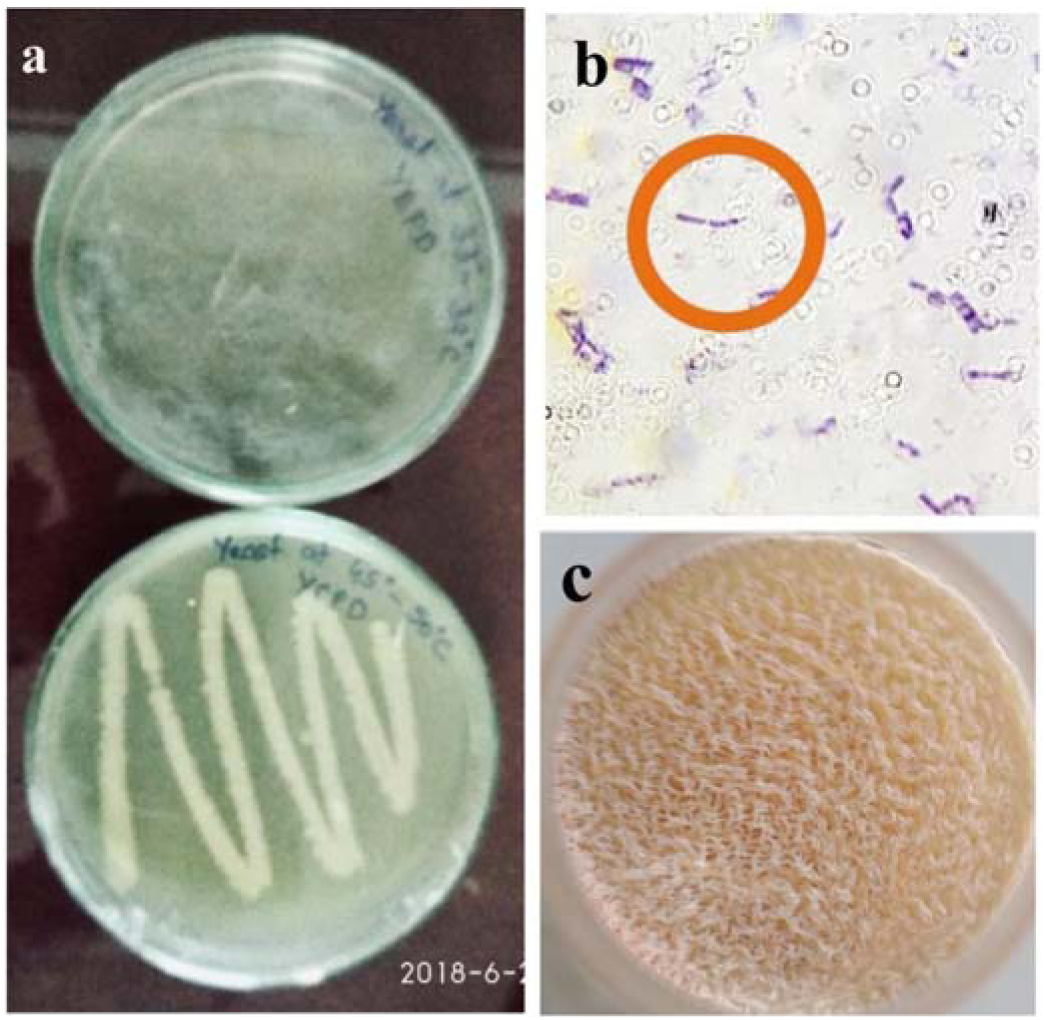
Isolated Strain TS-1. (a) Stain TS-1 on YePD agar plates one on the top is incubated at 33°-37°C and another is incubated at 45°-50°C. (b) Gram-positive rods. (c) A thick layer of Pellicle is formed on the YpSs broth when it is not incubated in a shaker.

### Synthesis and extraction of AgNps & EPS

The synthesis of AgNPs was easily noticed due to the colour change of the cell-free supernatant. As in the previous studies that involved silver nanoparticles production suggests the same where the supernatant with AgNO3 converts into brown colour on the synthesis of silver nanoparticles (Fig.2.b). While in control, the Cell-free supernatant remains in the light yellowish colour as same as broth colour (Fig.2.b). (Ahmed s. et al. 2015) (Guangquan Li. et al. 2012) (Deljou.A et al. 2016)

**Fig.2.**
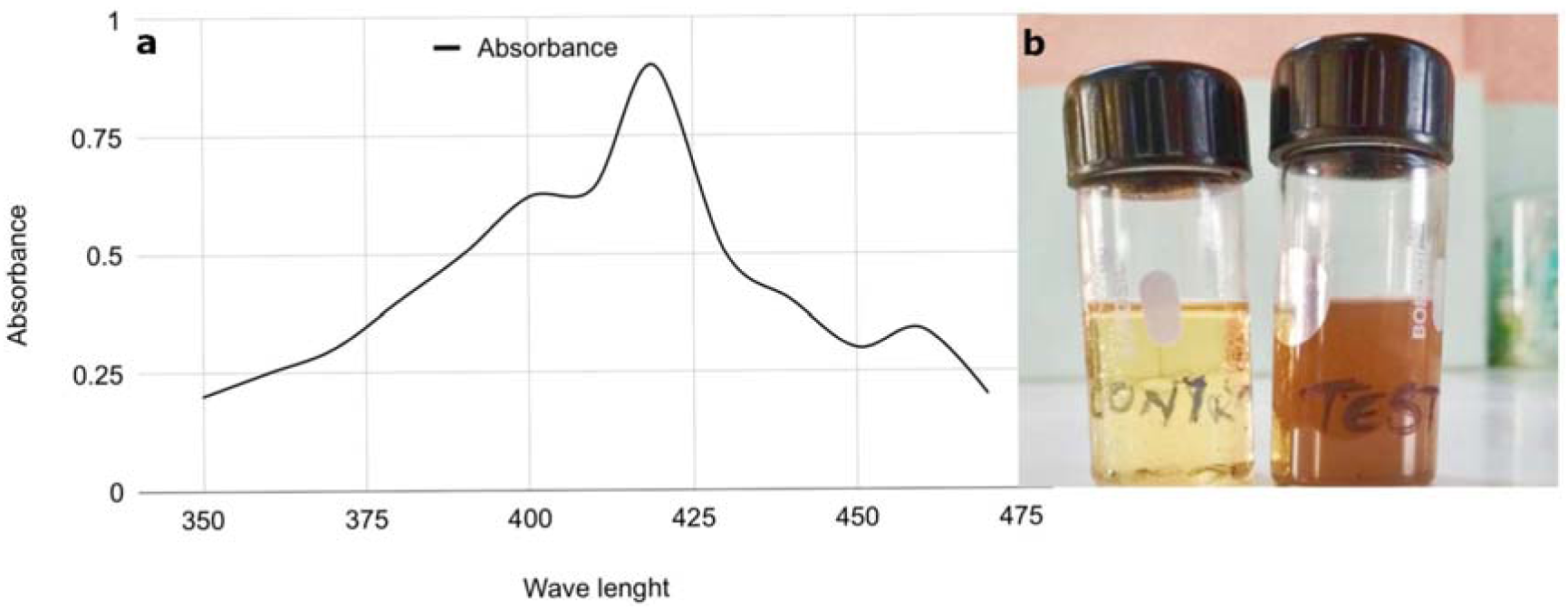
(a) UV Visible spectrum of AgNps. (b) visual colour changes from yellow to brown.

EPS can be observed as an opaque white substance due to precipitation after addition of absolute alcohol to the cell-free supernatant. EPS on drying it becomes light brown coloured powder.

### Characterization of AgNps & EPS

#### Uv-Vis Spectral analysis

From the addition of AgNO_3_ to the supernatant, the synthesis of silver nanoparticles begins where we can observe a peak at 419nm (Fig.2.a) which gradually strengthens till 24hrs that describes the Plasmon absorbance of synthesized AgNPs which is a direct confirmation of the nanoparticles synthesis. Silver nanoparticles synthesized using *Z. armatum* leaves extract also showed the absorbance peak at 419 nm (Jyoti et al. 2016). In the study carried out by (Rehab et al. 2019) Surface Plasmon resonance of the AgNPs and the blue shift was observed from 428 to 422 nm. Size, morphology, shape, composition, and dielectric environment of the synthesized nanoparticles influence the Surface plasmon absorbance band.

#### Scanning electron microscope (SEM)

It is evident that synthesized nanoparticles have a variation in particle sizes (Fig.3.d,e) and the average size estimated was 20nm visually compared with the scale given on the SEM images. The particles observed to be in the size range from 10 nm to 50 nm (Fig.3.c), which seemed to increase as the concentration of AgNO_3_ increased (Fig.3.c,d,e).

**Fig.3.**
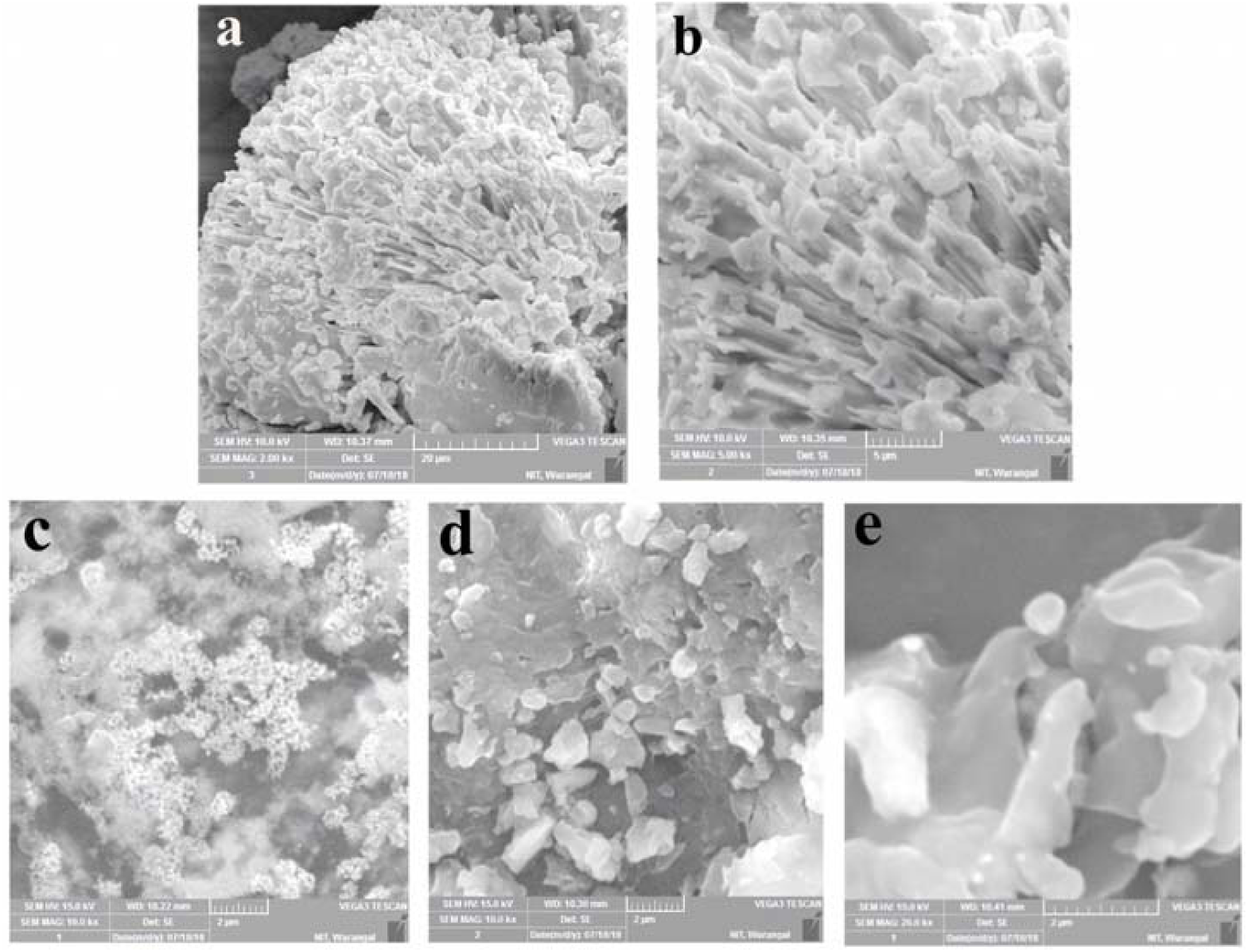
SEM images of (a,b)EPS & (c,d,e)Nanoparticles. (a) EPS at 2000x magnification (b) EPS at 5000x magnification (c) AgNps at 10000x magnification (1mM) (d) AgNps at 10000x magnification (5mM) (e) AgNps at 20000x magnification (10mM)

Surface morphology of EPS is characterized by SEM, which reveals a compact, porous and coarse surface. Upon 5000x magnification EPS from *Bacillus amyloliquefaciens* TS-1 was observed to have the weblike structure (Fig.3.a,b). Whereas the case of EPS produced from *Lactobacillus plantarum* also has a porous nature described by(Sri Laxmi et al. 2017). EPS extracted from *B. anthracis* is compact with a smooth surface, and without any porous features (Aparna et al. 2017) such differences in the topology and morphology are mainly caused by the difference in Physicochemical properties of the difference Exopolysaccharides. Extraction, purification and preparation techniques also play a role in changing the surface morphology of EPS. (Sri Laxmi et al., 2017)

#### Energy-Dispersive X-ray spectroscopy (EDAX)

The EDAX Spectrum from the (Fig.4.a) reveals the highest peak of elemental Ag at 3kev for the silver nanoparticles suspension, which directly suggests that silver is the predominant element in the respective product. The EDAX spectrum reported by (Singh et al. 2017) also shows the peak of Ag at 3kev for silver nanoparticles. The carbon peak at 0.3kev in the spectrum corresponds to the SEM grid utilized for the study.

**Fig.4.**
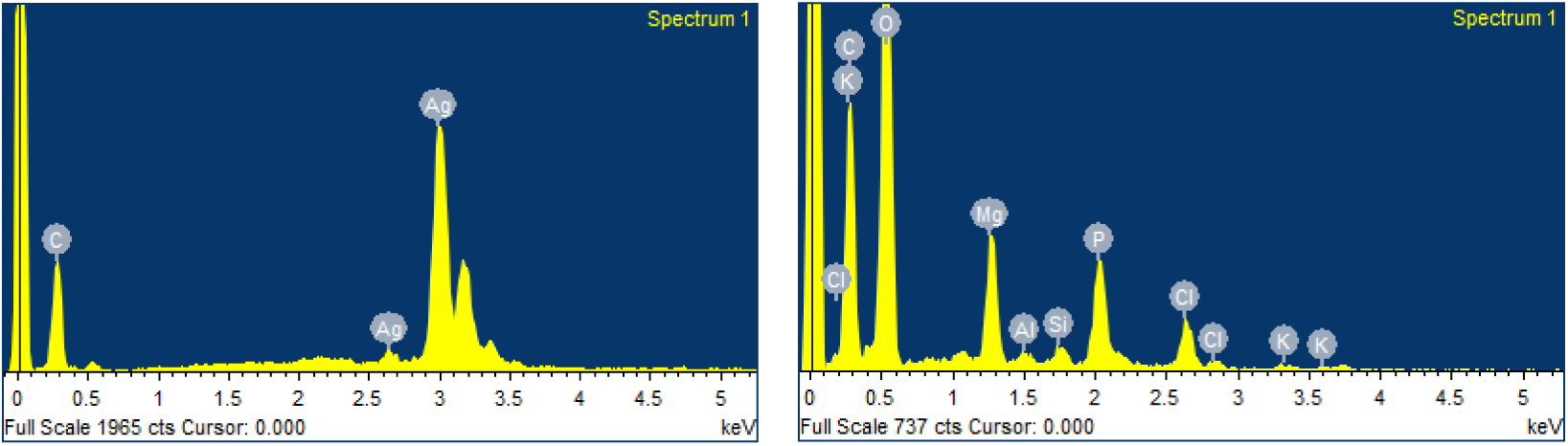
EDAX images. (a) Nanoparticles (b) EPS

**Fig.5.**
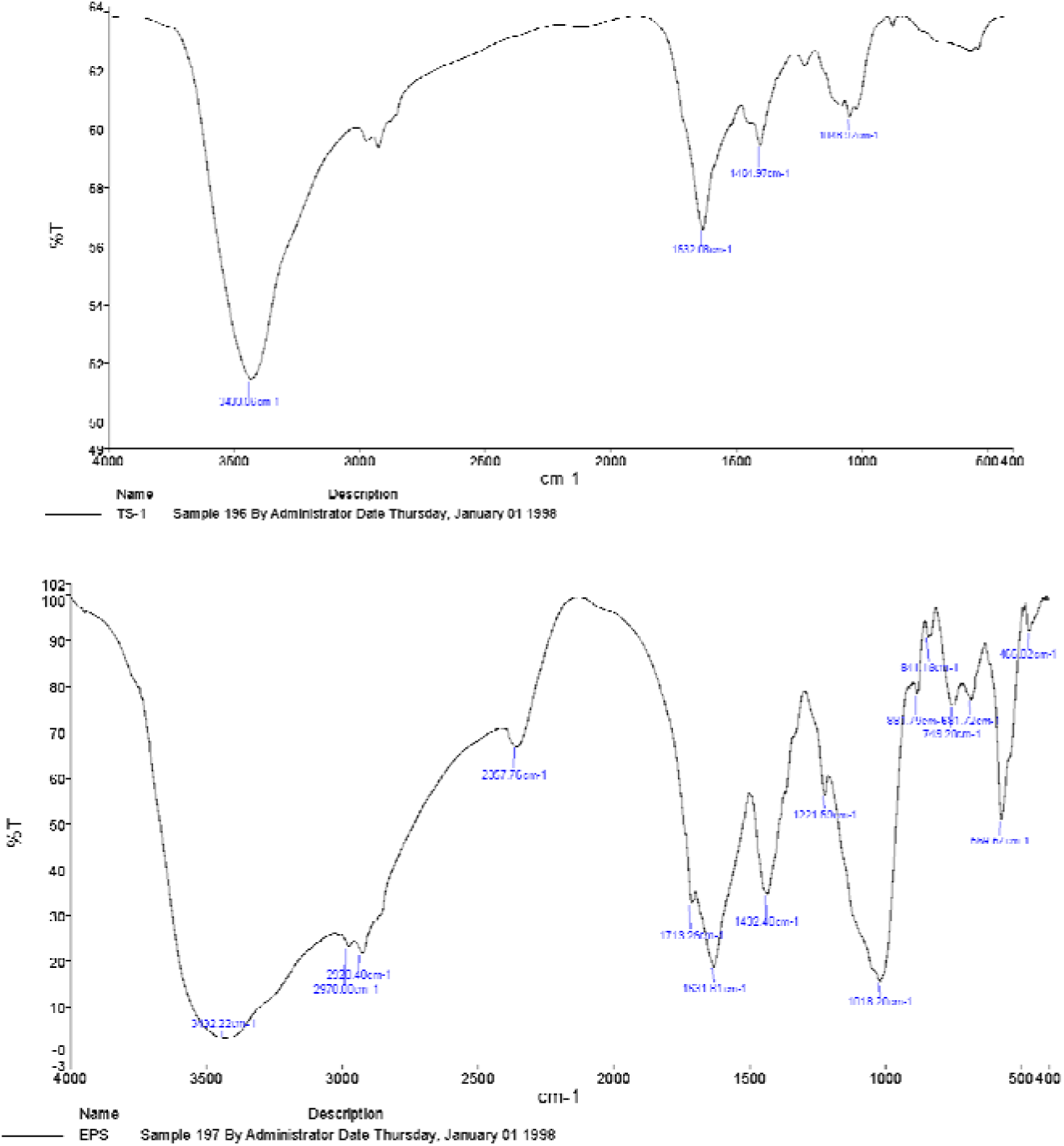
FTIR spectrum. (a) Nanoparticles (b) EPS

**Fig.6.**
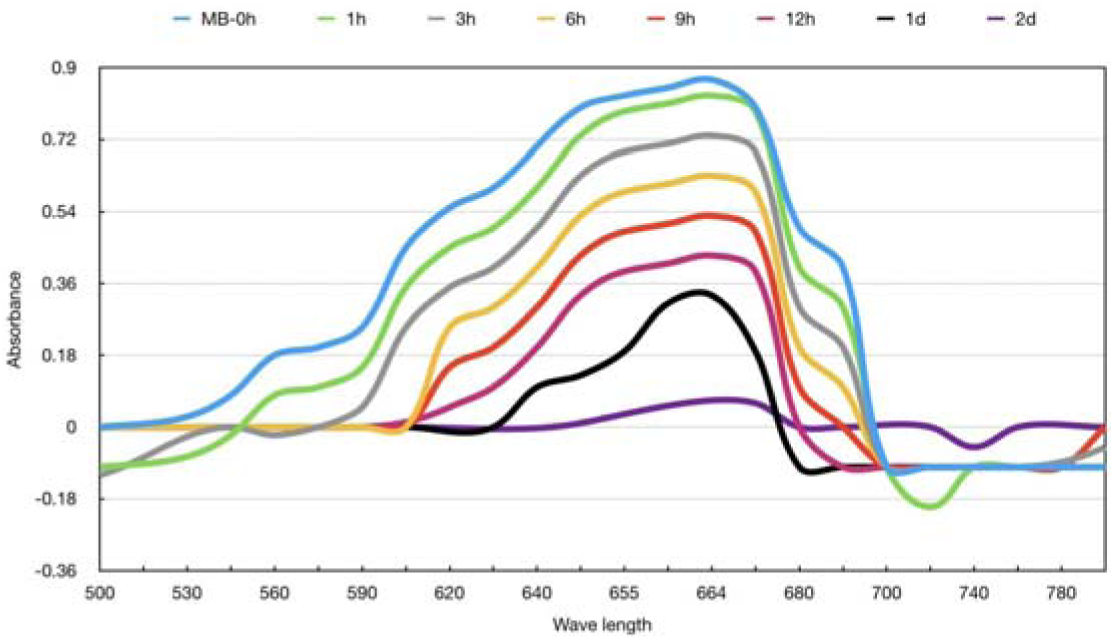
Uv Spectrum of Methylene blue for 48hrs of incubation with AgNPs.

The elemental composition of the EPS from the EDAX spectrum showed the composition of carbon, oxygen, magnesium, aluminum, silicon, phosphorus, potassium and chlorine. Based on our literature search, there were no reports showing the surface morphology and elemental composition of the EPS produced by *Bacillus amyloliquefaciens.* Spectrum also showed the presence of 41.23% of carbon, 48% of oxygen further confirmed the resultant polysaccharide compounds. The present result shown in the spectrum can be supported by the article showing presence of chlorine which is also reported in the EPS produced by *Streptococcus thermophilus* (Sri Laxmi et al. 2017). The presence of Carbon and Oxygen in high percentages is significantly a composition mainly consisting of sugar moieties. The amounts of C, H, N, and O in EPS extracted from *L. delbrueckii subsp.* were 39.1, 6.2, 2.8, and 50.3, respectively(Goh et al. 2005). (Wang et al. 2019) reported the amounts of C, H, N, and S in crude EPS extracted from *L. sakei* were 39.3, 6.1, 0.4, and 0.1, respectively.

#### Fourier transforms infrared (FTIR) spectroscopy

FTIR spectrum of nanoparticles describes the reducing & capping agents which are involved in stabilizing the nanoparticles. The spectrum showed a number of peaks, thus reflecting its complex nature. The strong, broad absorption band at 3433.06 cm-1 is characteristic of the alcohol/phenol —OH stretching vibration, carboxylic acid —OH stretch and N-H stretching of amides. The strong peak at 1632.08 cm-1 is characterized to —NH stretch of primary amines. The peaks between the ranges 1400-1000 cm-1 were assigned to the C-F stretching. These results suggested that the biological molecules possibly perform a dual function of synthesis and stabilization of silver nanoparticles (Singh et al. 2018) (Rehab et al. 2019). The present results of the spectrum can also be supported by the data reported by (Metuku et al. 2014), who showed the FTIR spectrum confirming the presence of protein-like compounds on the surface of synthesized silver nanoparticles. Later on, confirmed that these proteins act as the capping agents which maintain the stability of nanoparticles. Finally, this spectrum reports the presence of proteins in the synthesized nanoparticles.

FTIR spectrum from Fig.4.b. showing peaks at 3432.22 is a Medium N-H stretching of primary amine, 2928.48, 2978 are a Medium C-H stretching of alkane group, 2357.76 is Strong O=C=O stretching of carbon, 1713.26 is Strong C=O stretching of an aliphatic ketone, 1631.81 is Medium C=C stretching of disubstituted alkene, 1432.40 is Medium C-H bending of alkane group, 881, 841, 749 are strong peaks of the beta glycosidic bond. Thus the EPS is probably constituted of monosaccharide sugar units that are interlinked with each other using beta glycosidic bonds. (Aparna et al. 2017)

#### Degradation of Methylene blue

Degradation of azo dyes is important these days as these dyes are one of the main contaminants to the water. The absorption peaks of methylene blue dye in distilled water were observed at 664 nm in the visible region as shown in Fig-7 as MB-0h, which is also mentioned in (Jyoti et al. 2016), (Shahwan, T. et al. 2011). In the presence of AgNPs, the absorption spectrum showed dispersed peaks after the incubated time of 48 hours. This shows the degradation of methylene blue after the addition of silver nanoparticles which confirms the catalytic activity of AgNPs.

## Conclusion

It has been shown for the first time that the use of the thermophilic bacterium, *B.amyloliquefaciens* could be used for extracellular synthesis of metal (silver) nanoparticles and EPS. The stability of the nanoparticle solution could be due to the secretion of certain reducing enzymes and capping proteins by the bacterium. The advantages offered by using a bacterium mediated synthesis of metallic nanoparticles and exopolysaccharide include a toxin-free synthesis. The economic viability of the method is also the ease in handling large scale synthesis. While AgNPs produced by this bacterium also acts as a catalyst in degradation of Methylene blue dye.

## Abbreviations

AgNp’s: Silver Nanoparticles
EPS: Exopolysaccharide
Uv: Ultraviolet
EDAX: Energy Dispersive X-ray Spectroscopy
SEM: Scanning electron microscopy
FTIR: Fourier Transform Infrared Spectroscopy
NAM: Nutrient Agar Media
YpSs: Yeast powder Soluble Starch
YePD: Yeast extract Peptone Dextrose

## Acknowledgments

I sincerely acknowledge Dr. Koteshwar Rao, Dr. G.Krishna, Dr. Aparna Banerjee for their help in analyzing the data. Authors thankful to the Head of the Department of Biotechnology, Chaitanya Degree and PG College for their persistence support during the project. Our sincere thanks to Dr. P.Sreenivasa Rao for his guidance and support during this project.

## Compliance with ethical standards

The authors confirm that there are no conflicts of interest.

